# Fetal programming by sodium saccharin and damage on male offspring reproductive

**DOI:** 10.1101/2021.10.15.464538

**Authors:** Alana Rezende Godoi, Vanessa Caroline Fioravante, Beatriz Melo Santos, Francisco Eduardo Martinez, Patricia Fernanda Felipe Pinheiro

## Abstract

Male infertility is responsible for 20-70% of infertility in couples. We investigated the effects of fetal programming with sodium saccharin consumption in testis structure and function and in male offspring fertility. Feed intake and efficiency, organ and fat weight, quantification and expression of AR and PCNA proteins, sperm count and hormonal dosages were performed. Changes in consumption were found in the final weeks of the experiment. Decreases in the expression and quantification of AR and PCNA, tubular diameter and luminal volume, and increase in epithelial and interstitial relative volumes were observed. Lower sperm count and transit and lower estradiol concentration were also found. The consumption of sodium saccharin by the dams programmed the male offspring affecting the HPG axis with alterations in Sertoli cell proliferation, AR expression and quantification, and sperm count. We hypothesize that these changes may be due to the reduction of estradiol that caused the loosening of the tight junctions of the blood-testis-barrier (BTB), causing cell losses during spermatogenesis, also reflecting, under the decrease in tubular diameter with an increase in epithelial volume and consequent decrease in luminal volume. Sodium saccharin programming directly affected the reproductive parameters of male offspring and adult fertility.

## Introduction

Infertility is a global public health issue, as it affects a significant proportion of humanity (https://www.who.int/reproductivehealth/topics/infertility/perspective/en/, acess in 05/10/2021). Male infertility is responsible for 20-70% of the causes of infertility in couples and it is estimated that more than 30 million men worldwide are infertile. (AGARWAL et al., 2015).

There are male reproductive health disorders related to abnormal testicular development that can alter hormonal secretions and cause a malfunction in the Leydig or Sertoli cells (SKAKKEBAEK et al., 2001, 2007). Additionally, the prevalence of these pathologies has been associated with environmental and lifestyle factors, such as a sedentary lifestyle, exposure to toxic agents, and diet (SHARPE & SKAKKEBAEK, 2008; SKAKKEBAEK et al., 2001).

Reports of obesity have increased over the years, and from 1975 to 2020 the cases tripled (https://www.who.int/news-room/fact-sheets/detail/obesity-and-overweight, acess in 05/10/2021). It is estimated that by 2030, 58% of the world’s adult population will be overweight (KELLY et al., 2008). To help individuals in treatments for weight loss and diabetes control, the consumption of artificial sweeteners, sugar substitutes, has been growing exponentially and gaining popularity (CUMMINGS & OVERDUIN, 2007; POLYÁK et al., 2010). Of the non-nutritive sweeteners, sodium saccharin is one of the most consumed worldwide and is considered by the *Food and Drug Administration* (FDA) (https://www.fda.gov/food/food-additives-petitions/additional-information-about-high-intensity-sweeteners-permitted-use-food-united-states, acess in 05/10/2021) as safe because it is excreted completely unaltered by the kidneys (WHITEHOUSE, BOULLATA, MCCAULEY, 2008). However, its use is restricted during the gestational period, as there is placental permeability, which may interact with the conceptus and remain in fetal tissues due to its lower excretion capacity. It can be part of breast milk when ingested during lactation (BRUGNERA, BARUFFI, PANATTO, 2012).

Therefore, we must consider the hypothesis of “Developmental Origins of Health and Disease (DOHaD)” which treats the individual’s adaptive responses under specific nutritional conditions early in life. DOHaD is characterized by the organism’s limited susceptibility to a critical ontogenic window and the emergence of lasting effects in the body’s structure or function (MCMILLEN, ROBINSON, 2005; RINAUDO, WANG, 2012).

It is considered the existence of a “Masculinization Programming Window” (MPW), between 15.5-18.5 days post-conception, in the development of the male genital system of the rat. During this period, androgens act in the formation, proliferation, and normal growth of male organs and structures (WELSH et al., 2008; MACLEOD et al., 2010).

A recent study, involving the use of sodium saccharin in male mice, showed negative reproductive effects. Changes in LH levels were found, which consequently reflected in the production of testosterone. Furthermore, has been suggested that high levels of sodium saccharin may initiate the apoptotic cascade via caspase-3 through the binding of this food additive to bitter taste receptors (T2Rs). The effects of saccharin sodium led to changes in sperm quality including lower sperm count, sperm viability and motility, and increased sperm abnormalities (GONG et al. 2016).

Considering that the consumption of sodium saccharin can modify the intrauterine environment and damage reproductive health, we proposed to investigate the effects of fetal programming with the use of sodium saccharin during pregnancy and lactation in the structure and function of the testis and in the fertility of male offspring.

## Methods

### Animals

#### Mating

Sprague-Dawley rats 90-days-old were obtained from the State University of Campinas (UNICAMP, Campinas, São Paulo, Brazil). Males (n = 10) and females (n = 20) were housed in maternity boxes, in the ratio of 2 females to 1 male for mating. After confirmation of gestational, the dam rats were placed in individual boxes for the formation of experimental groups. Throughout the experimental period, the rats were kept in the *Bioterium* of the Department of Structural and Functional Biology (Anatomy Division) of the Biosciences Institute of Botucatu (IBB/UNESP, Botucatu, São Paulo, Brazil) under controlled conditions of temperature (22 ± 2 °C), humidity (50 ± 10%) and 12h light/dark cycle. The Ethics Committee on the Use of Animals of IBB/UNESP (number 1078 CEUA) approved this protocol.

#### Experiment

From gestational day 0, 10 dams received only food and water *ad libitum* (control group) while another 10 dams received food and water *ad libitum* and liquid diet sweetened with 0.3% sodium saccharin (Dinâmica^®^) (sodium saccharin group) during the pregnancy and lactation. On the day of birth, 4 males and 4 females were kept for equal food availability. After weaning, the male offspring of both groups received only food and water ad libitum. Euthanasia occurred at 120 days of age (Fig. 1).

**Figure 1.**
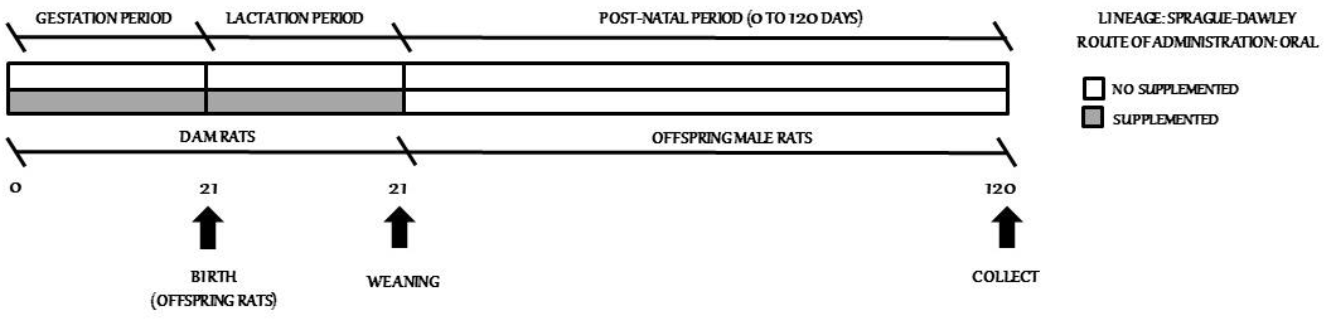
Experimental design.

#### Liquid diet

The liquid diet offered was prepared by adding 20 mL of low-fat plain yogurt (Nestlé^®^) to 15 mL of sodium saccharin solution (0.3%). A mixer was used to homogenize the diet. Only animals which ate at least 70% of the diet were included in the study. This experimental design was adapted of Cervantes-Rodríguez et al. (2014), Feijó et al. (2013), Pinto et al. (2017), and Swithers & Davidson (2008).

### Analysis

#### Body weight, food intake and feed efficiency

From weaning to the death of the animals, the body weight, water intake, and male offspring data were weekly measured. Feeding efficiency was validated by the food efficiency coefficient (FEC: (*Final weight – Initial weight/Feed consumed amount*) and weight gain for caloric consumption (*WGCC: (Final weight – Initial weight)/Kilocalories consumed amount*) (Campbell, 1963).

#### Collect of biological material

The animals fasted for 6 h before collection. Subsequently, the rats were weighed and anesthetized with sodium thiopental (40 mg/kg) and killed by decapitation. After blood collection, an abdominal-pelvic laparotomy was performed, and the testes and fats were dissected and weighed.

#### Determination of the adiposity index

The fats on the epididymis, cardiac, visceral and retroperitoneal fats were collected and the absolute weights were measured. The adiposity index was determined by (fats epididymis + cardiac + visceral + retroperitoneal/final body weight) ^*^100.

#### Light Microscopy

The testes were fixed in a 10 % formaldehyde solution in sodium phosphate buffer (PBS) (pH 7.4), washed for 48 h in running water and placed in a 70 % alcohol solution.

Subsequently, they were dehydrated in growing solutions of alcohols (ethanol and butyl), included in paraplastic and cut to 4 μm thick. Collections of slides stained with hematoxylin-eosin (HE) were obtained. The slides were analyzed and photographed using a BX 41 -2 photomicroscope with a digital camera, model SIS-SC30, Olympus from the Department of Structural and Functional Biology (Anatomy Division) of the IBB/UNESP.

#### Immunohistochemistry

Histological sections of the testes were placed on silanized slides and remained for 1 hour in an oven regulated at 58 °C. Then, the histological sections were deparaffinized in xylol and hydrated in decreasing alcohol (ethanol) solutions, running water for 10 minutes and in PBS buffer (pH 7.4). Subsequently, the slides in 0.01 M sodium citrate buffer (pH 6.0) were subjected to microwave oven heat (700-800 W) during 15 minutes, divided into 3-5 minute cycles. The slides which were incubated with anti-androgen receptor antibody were placed in 0.01 M sodium citrate buffer (pH 6.0) in a pressure cooker and subjected to high pressure and temperature for 60 minutes. Then, the slides were placed in a 3 % hydrogen peroxide solution in methanol during 15 minutes. To block nonspecific reactions, the slides were incubated in a 2 % skimmed milk solution (Molico^®^) in PBS buffer (pH 7.4) during 1 h. In the next step, the histological sections were subjected to reaction with Proliferating cell nuclear antigen (PCNA-PC-10 Novocastra^®^, dilution 1: 100 in PBS, pH 7.4) and Androgen receptor (AR -N-20 Santa Cruz Biotechnology Inc.^®^, dilution 1: 100 in PBS, pH 7.4) and incubated in a humid chamber overnight at 4 °C in a refrigerator. After incubation, the slides were washed in PBS buffer (pH 7.4) and the sections were incubated for 1 h at room temperature with biotinylated IgG anti-mouse secondary antibody which is produced in rabbit (E 0354 -Dako Cyt. Inc.^®^, 1: 100 dilution in PBS, pH 7.4) and biotinylated anti-rabbit IgG antibody which is produced in goats (E 0432-1 Dako Cyt. Inc.^®^, 1: 100 dilution in PBS, pH 7.4). After this step, the slides were washed with PBS (pH 7.4) and submitted to the avidin-biotin-peroxidase solution (ABC Kit -PK -6100 Vector Laboratories®) for 45 minutes. Then, the slides were washed in PBS (pH 7.4) and subjected to diaminobenzidine (DAB) (di-amino-benzidine-Sigma^®^) for 5 minutes. Subsequently, the slides were washed in tap water and contrasted with Harris’ hematoxylin. During the technique, positive and negative controls were obtained. The slides were analyzed and photographed using a BX 41 -2 photomicroscope with a digital camera, SIS-SC30 model, Olympus from the Department of Structural and Functional Biology (Anatomy Division) of the IBB/UNESP.

### Morphometric and stereological analysis

The PCNA^+^ sperm and Sertoli AR^+^ cells were measured. To determine the indexes (total number of positive cells/seminiferous tubules), 180 tubules per group were evaluated. The described variables were analyzed using the image analysis program Cellsens Standard (Olympus) from the Department of Structural and Functional Biology (Anatomy Division) of the IBB/UNESP.

The diameter of the seminiferous tubules was evaluated in three histological sections of each animal.

For the stereological evaluation of the testis, 100 histological fields of each animal (5 animals/group) were analyzed. The images were evaluated using the method described by Weibel (1978) using a 168-point grid.

### Protein extraction and Western Blot

The testes were homogenized in extraction buffer containing, Triton-x-1 %, 150 mM NaCl, 10 mM Tris pH 7.4, 1 mM EDTA, 1 mM Hepes, pH 7.6, 0.2 mM PMSF and 10 µL/mL of protease inhibitor cocktail. Testicular extracts were obtained by centrifugation for 20 minutes at 4000 rpm at 4 °C. An aliquot of each sample was used to determine the protein concentration, using Bradford’s reagent. The corresponding 60 µg of protein was applied to the SDS-polyacrylamide gel. After electrophoresis, the material was transferred electrically to nitrocellulose membranes. The membranes were then blocked with 3 % skimmed milk (Molico^®^) diluted in TBS-T for one hour and incubated with the primary AR antibodies (AR N-20 Santa Cruz Biotechnology Inc.^®^, 1: 100 dilution in skimmed milk 1 %) and PCNA (PC-10 -VP-P980 Vector Laboratories^®^ and PC-10 Novocastra®, dilution 1: 500 in skim milk 1 %). After washing with TBS-T buffer, the membranes were incubated for 2 h with secondary anti-rabbit IgG peroxidase antibodies, antibody produced in goats (A0545 -Sigma^®^, 1: 20.000 dilution in 1 % skimmed milk) and anti-mouse IgG peroxidase, antibody produced in goats (A9044 -Sigma^®^, 1: 20.000 dilution in 1 % skimmed milk). After a new series of washes with TBS-T, the bands were developed with an Amersham ECL Prime Western Blot Detection Reagent (RPN2236 -Amersham^®^) chemiluminescent substrate on the G-BOX equipment. B-actin was used as an endogenous control. The intensity of the staining obtained on the different targets was determined through pixel counting using the Image J^®^ software (version 1.48, Rasband, W.S., Image J, U. S. National Institutes of Health, Bethesda, Maryland, USA, http://imagej.nih.gov/ij/, 1997-2014).

### Daily testicular production

The testicular parenchyma was thawed, placed in a 15 mL test tube and weighed, adding 5 mL of STM solution containing 0.9 % NaCl and 0.05 % Triton X 100 and homogenized. A new mixture of the homogenate was obtained by the dilution with STM in the proportion of 1:10. Subsequently, a sample was transferred to two Neubauer chambers, divided into 2 antimers and the spermatids in stage 19 were counted in 5 fields per antimer. The number of spermatids was obtained by the average of the counts multiplied by the dilution factors. In the sperm concentration (number of sperm/g of testis), the average number of sperm was divided by the weight of the testicular parenchyma. The calculation of daily sperm production consisted of dividing the sperm quantity by 6.1 (a factor that corresponds to the number of days in which mature spermatids, stage 19 of spermatogenesis are present in the germinal epithelium).

### Counting the number and transit time of sperm

The collected epididymis was divided into the head (H) + body (B) and tail (T), weighed and frozen. When evaluating the transit time of sperm in the epididymis, the H + B and T segments were prepared as follows: for each 200 mg of H + B, 1mL of STM solution was added and for each 100 mg of T, 1mL of STM. Subsequently, the samples were submitted to the homogenizer. A new mixture of the homogenate was obtained by dilution with STM in the proportion of 1:20. The number of sperm in each segment was counted in two Neubauer chambers and 5 fields per antimere were counted. The transit time of sperm through the epididymis was calculated by dividing the number of spermatozoa by the value obtained in the daily production of sperm from each animal.

### Hormonal Analysis

Plasma concentrations of testosterone (ng/mL) and 17 β-estradiol (pg/mL) were determined by the competitive ELISA method. The ELISA kit: testosterone (Elabscience Biotechnology Co., Houston, Texas, USA, catalog n° E-EL-0155, 96 T, sensitivity of 0.17 ng / mL, detection range between 0.31-20 ng / mL, repeatability with coefficient of variation < 10 %, specificity proven with no significant cross-reaction or interference between T and analogs); estradiol (Elabscience Biotechnology Co., Houston, Texas, USA, catalog n° E-EL-0065, 96T, sensitivity of 25 pg / mL, detection range between 40-1500 pg / mL, repeatability with intra and inter coefficient of variation ≤15 %, proven specificity with no significant cross-reactivity or interference with progesterone and estriol). The samples were read on a spectrophotometer at 450 nm.

### Statistical analysis

The variables were analyzed using the GraphPad Prism program (version 8, GraphPad Software Inc., San Diego, CA). Firstly, the data were subjected to the Shapiro-Wilk normality and variability test. For parametric data, the variables were studied using t-test and analysis of variance (ANOVA) complemented with the Sidak multiple comparison test. For non-parametric data, the Mann-Whitney test was used. The results were expressed as mean ± SD (standard deviation) considering the level of significance of 0.05 %.

## Results

### The use of maternal sodium saccharin increased feed and water consumption of male offspring

Bodyweight and feed efficiency did not differ between groups (Fig. 2A e D), however, water and feed consumption was higher in the final weeks of the experiment (Fig. 2B e C).

**Figure 2.**
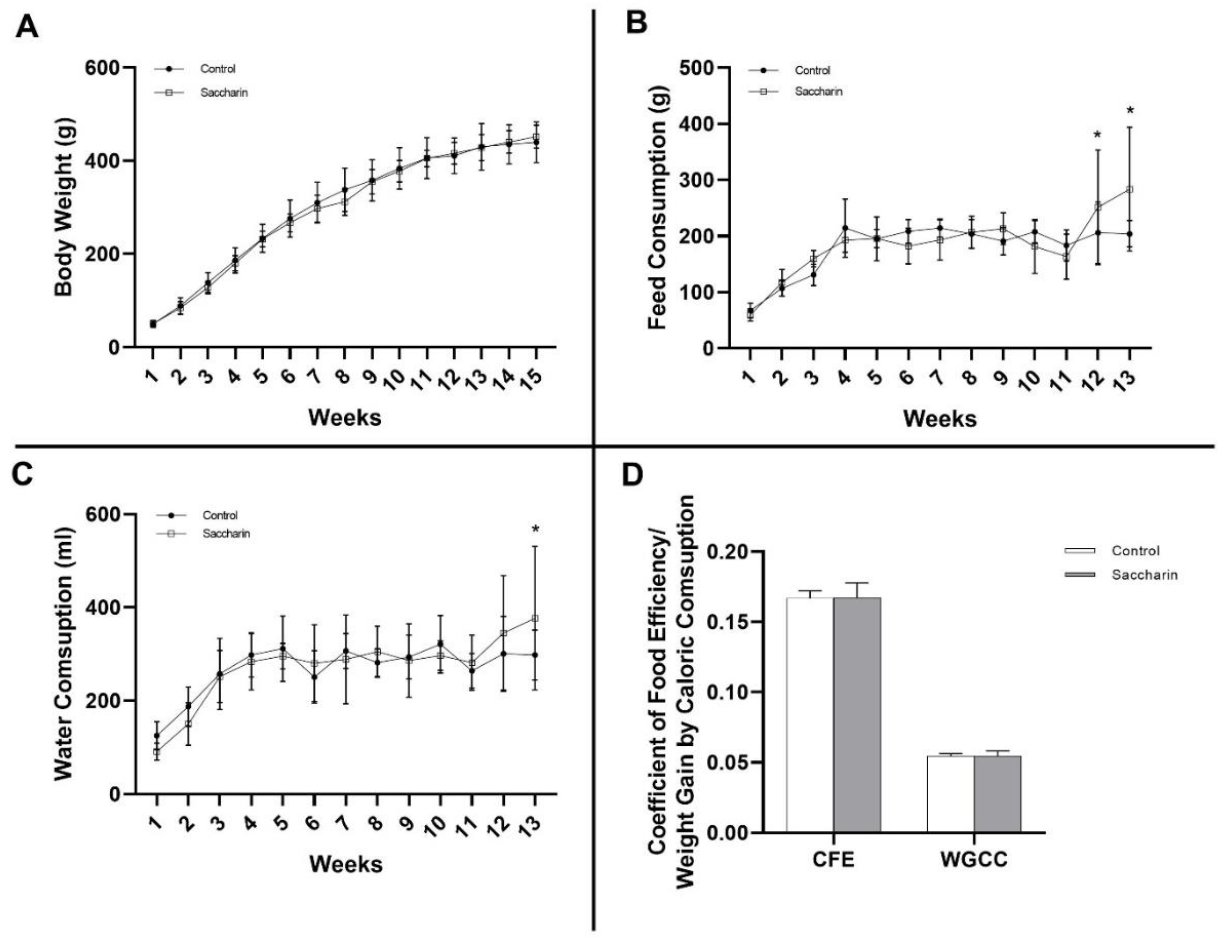
Parameters of male offspring from Control and Saccharin (n=12/group). A) Body weight, B) Feed consumption, C) Water consumption and D) Feeding efficiency of the male descendants. Values expressed as mean±SD. P values were calculated using a two-way ANOVA and a t-test. ^*^Significant differences between groups (p < 0.05) from post-hoc Sidak’s multiple comparison test. Figures 2A: P_Inter_= 0.9507, P_Week_= <0.0001, P_Group_= 0.2463; 2B: P_Inter_= <0.0001, P_Week_= <0.0001, P_Group_= 0.2240; 2C: P_Inter_= 0.0910, P_Week_= <0.0001, P_Group_= 0.7041.

### The use of maternal sodium saccharin did not change the organ weight and body fat of male offspring

There were no differences between the groups in the testis, epididymal, retroperitoneal, visceral, and cardiac weights, and adiposity index (Table 1).

**Table 1:**
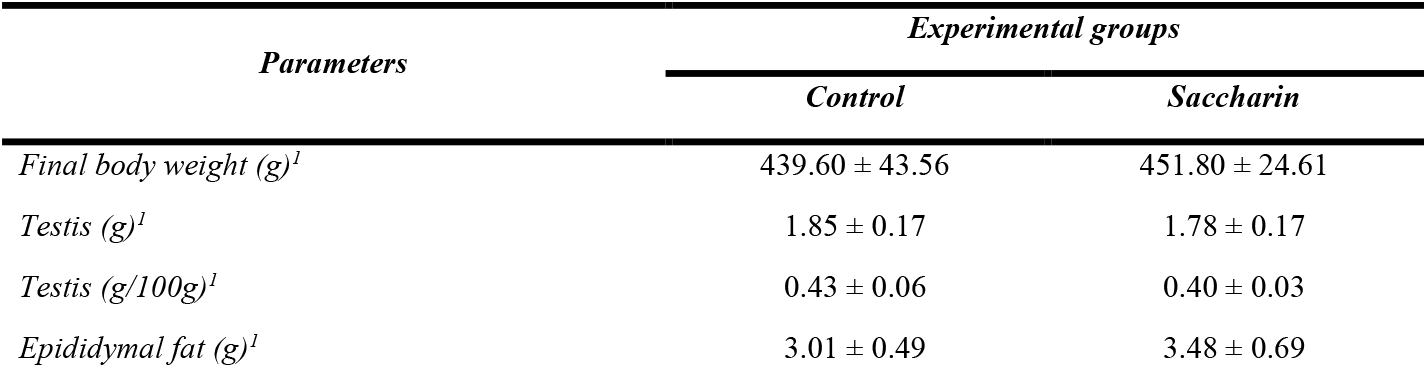
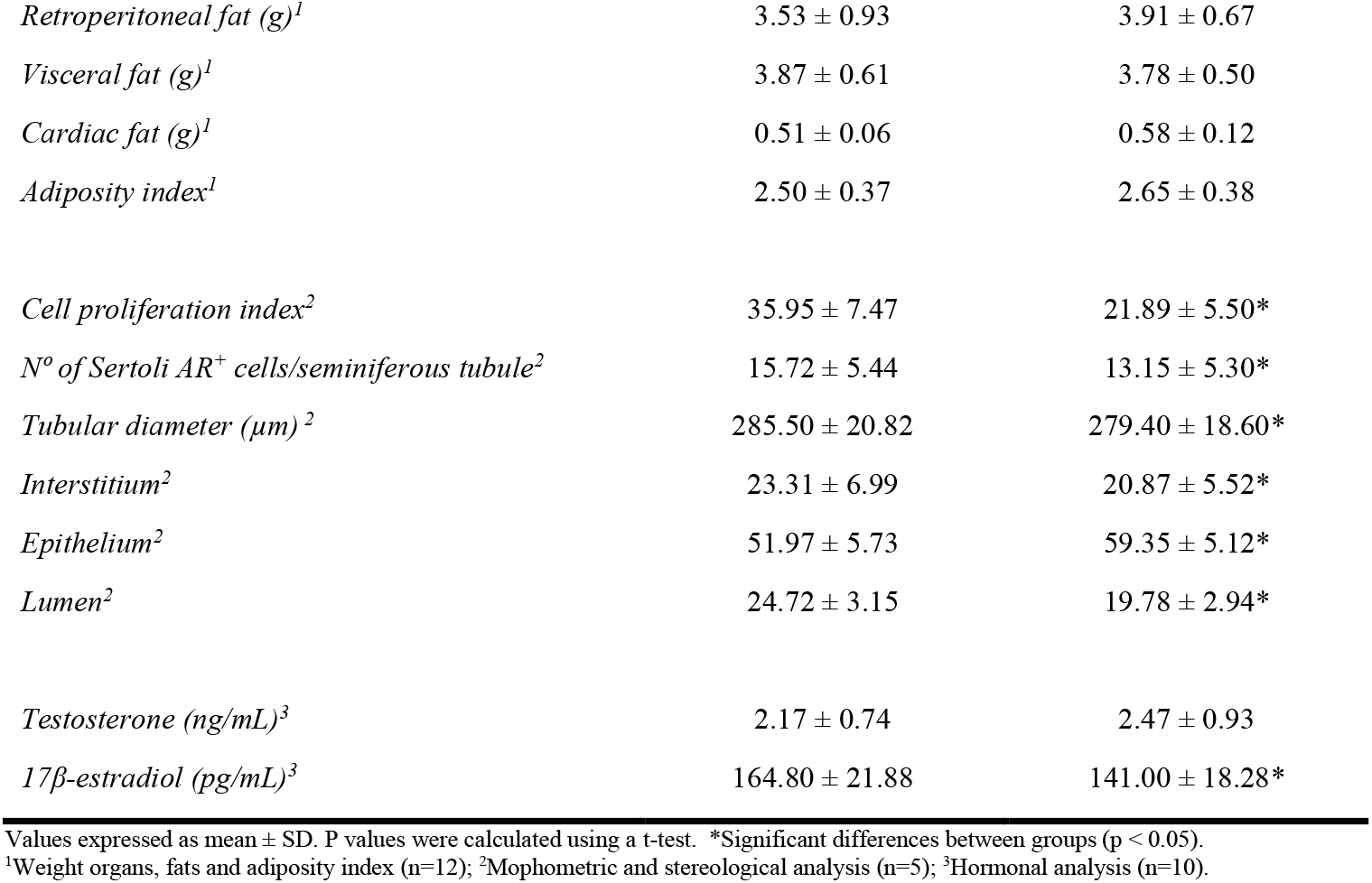
Body weight, absolute and relative weights of the testis, fat and adiposity index, cell proliferation index (number of sperm PCNA+/seminiferous tubule), the number of Sertoli AR+ cells/seminiferous tubule, tubular diameter (µm), the plasma levels of testosterone and 17β-estradiol, and the stereological analysis of male offspring from Control and Saccharin.

### The use of maternal sodium saccharin impaired the functionality of the male offsprings testis

The sodium saccharin group showed tubular morphological changes characterized by the presence of spermatocytes, elongated spermatids, and residual bodies in the tubular lumen (Figure 3 B e F). A decrease in the tubular diameter and in the relative volumes of the interstitium and lumen were also observed, in addition to an increase in the relative volume of the epithelium (Table 1).

**Figure 3.**
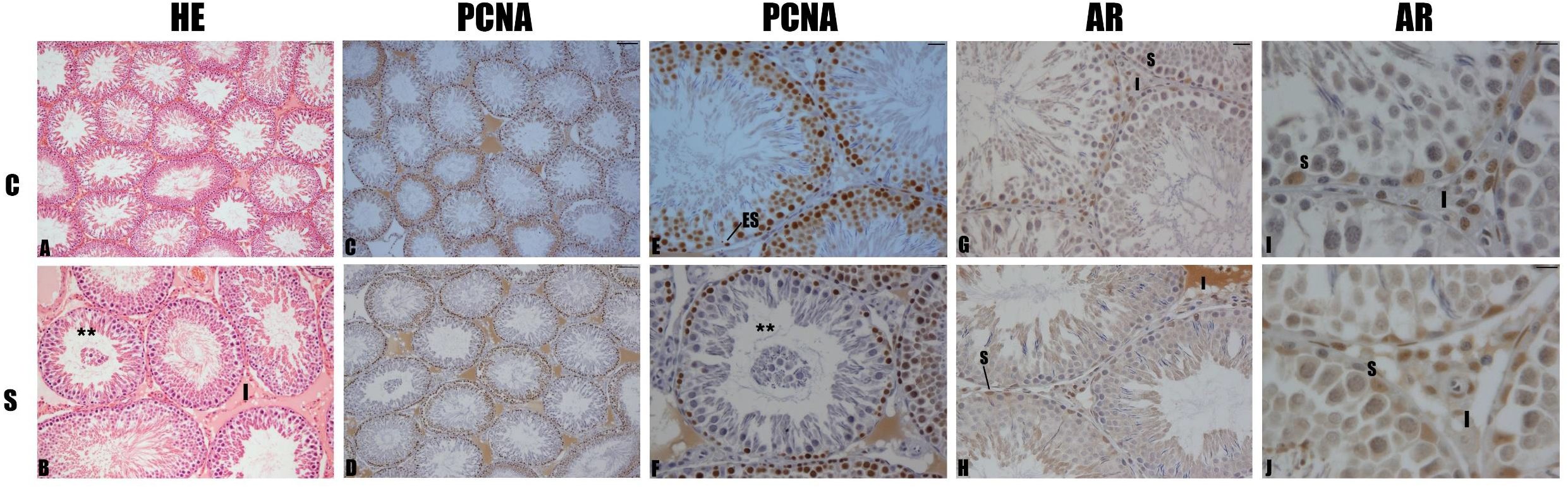
Cross sections of seminiferous tubules. C (Control) and S (Saccharin) groups (120 days old) (n=5/group). HE (Hematoxilin and Eosin); staining positive PCNA and AR. Spermatogonia (ES), Sertoli Cell (S), Leydig Cell (L), Interstitium (I), Seminiferous tubules with cells desquamated (^**^). (A, C and D) Barra = 100 μm, (B, E-F) Barra = 20 μm, (G-J) Barra = 10 μm.

The sodium saccharin group showed a decrease in cell proliferation indices and in the number of Sertoli AR+ cells. Quantitative analysis of PCNA and AR in the testis also showed decreased in the sodium saccharin group (Fig. 4).

**Figure 4.**
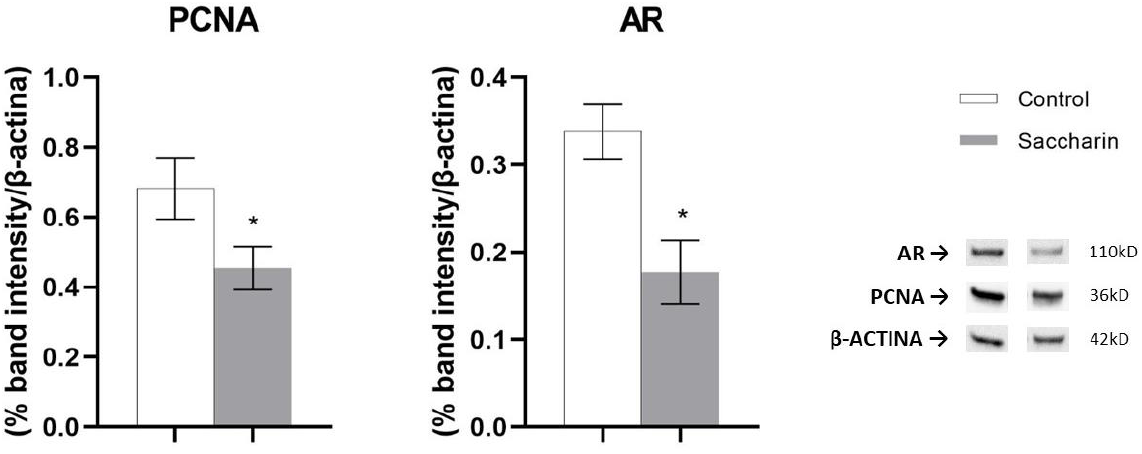
Optical densitometric analysis of PCNA and AR of the descendants from Control and Saccharin (n = 4/group). Values expressed as mean ± SD. Data analyzed by t-test. ^*^Significant differences between groups (p < 0.05).

The plasma concentration of 17β-estradiol was lower in the sodium saccharin group, while the plasma testosterone concentration showed no difference between the groups (Table 1).

### The use of maternal sodium saccharin impaired the reproductive efficiency of male offspring

The sodium saccharin group showed decreased sperm count and less transit in the tail region of the epididymis. The total sperm transit time was also shorter in the sodium saccharin group (Table 2).

**Table 2:** Sperm count in the testis and epididymis, transit time and sperm morphology of male offspring from Control and Saccharin.

| *Parameters* | *Experimental groups* |  |
| --- | --- | --- |
|  | *Control (n = 12)* | *Saccharin (n = 12)* |
| *Sperm count in the testis* |  |  |
| *Daily sperm production (x10<sup>6</sup>/testis/day)* | 13.77 ± 1.85 | 15.67 ± 2.34 |
| *Number of sperm cells in the testis (x10<sup>6</sup>)* | 83.97 ± 11.27 | 95.58 ± 14.25 |
| *Sperm count in the epididymis (head/body)* |  |  |
| *Number of sperm (x10<sup>6</sup>)* | 74.39 ± 16.75 | 83.23 ± 11.85 |
| *Sperm transit time (days)* | 5.42 ± 1.05 | 5.50 ± 1.14 |
| *Sperm count in the epididymis (tail)* |  |  |
| *Number of sperm (x10<sup>6</sup>)* | 245.10 ± 47.08 | 161.10 ± 36.34* |
| *Sperm transit time (days)* | 17.95 ± 3.38 | 10.24 ± 1.84* |
| *Total sperm transit time (days)* | 23.37 ± 4.20 | 15.74 ± 2.04* |

Values expressed as mean ± SD. P values were calculated using a t-test. \*Significant differences between groups (p < 0.05).

## Discussion

Environmental and nutritional insults during critical periods of fetal development bring permanent physiological or metabolic changes in the body, molding the body will respond to later challenges (Langley-Evans, 2009). Thus, we verified the morpho-functional response of the testis and the fertility potential of male offspring exposed to sodium saccharin during pregnancy and lactation.

The consumption of non-nutritive sweeteners stimulates compensatory food intake in response to the absence of calories related to the perception of sweet taste through the action of oral and intestinal sweet taste receptors that, through afferent vagal impulses, carry information to the thalamus and the system of reward (Bellisle & Drewnowski, 2007; Berthoud, 2002; Cummings & Overduin, 2007; Renwick & Molinary, 2010; Saris, 2003; Smeets et al., 2010; Yang, 2010). The absence of differences in weight gain observed in our study may be associated with the short period of exposure to maternal sodium saccharin consumption and the age of the animals.

The weight of the organs of the male genital system is a parameter for circulating testosterone levels (Clegg et al., 2001). Hormonal changes did not directly influence the absolute and relative weights of organs and fats. Although we did not observe differences in plasma testosterone, a decrease in AR expression and quantification of Sertoli AR+ cells were found. The role played by Sertoli cells in adult life is a result of the maturation process from birth to puberty. Testicular maturation is related to the maturation of the hypothalamic-pituitary-gonad (HPG) axis which, through negative and positive feedbacks, ensure adequate gonadal development and function. Androgens participate in several important processes for the maintenance of spermatogenesis, such as regulation of Sertoli cell maturation, Sertoli-Sertoli and Sertoli-germ cell junction involved in blood-testis-barrier (BTB) formation and maintenance, germ cell proliferation and differentiation (Wang et al., 2009; Ruwanpura et al., 2010; Verhoeven et al., 2010) and spermiation (O’Donnell et al., 2011).

During the testicular maturation phase, Sertoli cells respond to FSH (Orth, 1984; Al-Attar et al., 1997; Migrenne et al., 2012; Lambert & Bougneres, 2016; Kumar, 2018; Urrutia et al., 2019), other growth factors (Pitetti et al., 2013; Meroni et al., 2019; Neirijinck et al., 2019) and proliferate. This phase will have a direct influence on adult life because each Sertoli cell is capable of supporting a fixed number of germ cells. The amount of Sertoli cells produced during testis maturation will determine the sperm production in the adult (Orth, 1982 e 1984; Russell et al., 1990; Cooke et al., 1991; Hess et al., 1993).

Considering the MPW and changes in the intrauterine environment, such as stress or nutritional factors which can affect the development of the offspring, the consumption of sodium saccharin by dams during the gestational and lactational periods programmed the male offspring directly affecting the HPG axis and possibly influencing Sertoli cell proliferation during the testicular maturation period and in the expression and quantification of its androgen receptors, essential for the maintenance of spermatogenesis. In addition, the Sertoli cells are responsible for detecting developing sperm and supporting them until they mature and reach the apical region of the tubules, requiring the control of the BTB that separates and reforms the tight junctions through the N-cadherins. FSH combined with estradiol has been necessary for the transcription of N-cadherin mRNA, the protein responsible for cell-to-cell junction (MacCalman et al., 1994 e 1997). We verified cell losses in the lumen of the seminiferous tubules of animals programmed by sodium saccharin. These alterations may be due to the reduction of estradiol, which leads to the loosening of the tight junctions of the BTB, which can cause cell losses during spermatogenesis.

The cells responsible for the aromatization of testosterone to estradiol are Sertoli cells in the neonatal period, stimulating precursor germ cells through ER located in the plasma membrane (Zhou et al., 2002). When cells multiply and are capable of producing estradiol, positive paracrine feedback occurs and the hormone produced inhibits aromatases present in Sertoli cells (Boitani et al., 1981). Although estradiol is able to inhibit Sertoli cells, there is stimulation in germ cells. New evidence has shown that germ cells are responsible for the production of 50-60% of estradiol in the testes (Carreau & Hess, 2010; Carreau et al., 2012). In addition, estradiol and Platelet Derived Growth Factor (PDGF) has been shown to stimulate germ cell proliferation by giving Sertoli cells the regulation of mitosis through estradiol (Li et al., 1997; Thuillier et al., 2012). Cellular losses from the seminiferous tubules, therefore, may be helping to significantly reduce estradiol and, consequently, maintain cell proliferation with lower sperm counts in our results.

In addition, aromatase was observed in elongated spermatids and in human ejaculated sperm with a link between estradiol, capacitance and acrosome reaction. In a non-capacitating medium, only estradiol and aromatizable steroids were able to increase sperm mobility and migration (Aquila et al., 2003), validating the decrease in sperm transit in males exposed to sodium saccharin. We infer that male offspring programmed during fetal and neonatal development under the influence of sodium saccharin fail to convert testosterone to estradiol by aromatase. Although several studies demonstrate that the excess of intra-testicular estradiol causes damage to spermatogenesis (Schlegel, 2012), new findings show that the decrease in this hormone can also severely affect testicular morpho functionality (Schulster et al., 2016).

The stereological data of the animal’s testes programmed with sodium saccharin showed a smaller tubular diameter, accompanied by a larger epithelial volume and smaller luminal volume of the seminiferous tubules. Given the lower values found in the quantification of Sertoli cells in these animals, in addition to cell losses being found in the histological analyses, we assume that this decrease in tubular diameter is a reflection of the smaller number of cells present in these tubules. This, in turn, leads to an increase in epithelial volume and a consequent decrease in luminal volume.

In conclusion, maternal sodium saccharin consumption during pregnancy and lactation affects the reproductive parameters of male offspring, impacting spermatogenesis and fertile capacity in adulthood. Our study is a pioneer in fetal programming using sodium saccharin, a sweetener with growing use worldwide. Despite the limitations, we emphasize the risk of sodium saccharin on the reproduction of male offspring, highlighting the need for preventive measures and awareness of its use during pregnancy and lactation. Future studies may investigate whether these changes can be transmitted transgenerationally, characterizing development programming.

## Author contribution statement

ARG conceived the study, performed experiments, analyzed and finished data and wrote the paper. VCF and BML performed experiments. LAJJ, FEM and PFFP provided training to perform the experiments and intellectual input for the experimental design and data analysis. All authors contributed to edit the paper.

